# Estimating Ancestral States of Complex Characters: A Case Study on the Evolution of Feathers

**DOI:** 10.1101/2024.08.23.609354

**Authors:** Pierre Cockx, Michael J. Benton, Joseph N. Keating

## Abstract

Feathers are a key novelty underpinning the evolutionary success of birds, yet the origin of feathers remains poorly understood. Debates about feather evolution hinge upon whether filamentous integument has evolved once or multiple time independently on the lineage leading to modern birds. These contradictory results stem from subjective methodological differences in statistical ancestral state estimates. Here we conduct a comprehensive comparison of ancestral state estimation methodologies applied to stem-group birds, testing the role of outgroup inclusion, tree time scaling method, model choice and character coding strategy. Models are compared based on their Akaike Information Criteria (AIC), mutual information, as well as the uncertainty of marginal ancestral state estimates. Our results demonstrate that ancestral state estimates of stem-bird integument are strongly influenced by tree time scaling method, outgroup selection and model choice, while character coding strategy seems to have less effect on the ancestral estimates produced. We identify the best fitting models using AIC scores and a leave-one-out cross-validation approach (LOOCV). Our analyses broadly support the independent origin of filamentous integument in dinosaurs and pterosaurs and support a younger evolutionary origin of feathers than has been suggested previously. More generally, our study highlights that special care must be taken in selecting the outgroup, tree and model when conducting ASE analyses. With respect to model selection, our results suggest that considering a LOOCV approach, may yield more reliable results than simply comparing AIC scores when the analyses involve a limited number of taxa.

---

Feathers are a key novelty underpinning the evolution of avian flight. As such, knowledge of the sequence and timing of feather evolution is essential for understanding the origin of the avian body-plan. Exceptionally preserved stem-avians from the Late Jurassic - Early Cretaceous provide unparalleled insight into the evolutionary origin of feathers. Filamentous integuments with diverse morphologies have been documented in numerous dinosaur taxa (e.g. Xu 2020; O’Connor 2020) and even in pterosaurs (e.g. Kellner et al. 2010; Yang et al. 2019; Cincotta et al. 2022). These morphotypes range from simple structures to more complex ones and are associated with specific body regions and functions. Crucially, they are non-randomly distributed across the avian stem; the simplest morphotypes are widespread while more complex ones (such as pennaceous feathers) are restricted to taxa more closely related to the crown-group (Benton et al. 2019).

To date, at least 13 distinct morphotypes of feathers have been differentiated among integumentary structures reported from non-avian dinosaurs. Two basic categories of feathers are commonly recognized based on the general feather structure: pennaceous and plumulaceous types. Pennaceous feathers are characterized by barbs fused to a central shaft called the “rachis”, thus defining a clear planar surface (the vane). Plumulaceous feathers have no vane and are characterized by a rudimentary rachis and a tuft of barbs. Beyond this simple distinction, feather morphotypes can be further categorized according to various criteria including number of filaments (single or multiple), general morphology, secondary branching (e.g. barbules allowing interlocking of the barbs through hooklets), and symmetry of the vanes. This leads to the following categories: monofilaments, basally-joined filamentous feathers, filamentous feathers with a central filament, pennaceous feathers, and asymmetrical pennaceous feathers.

Testing hypotheses about the timing, sequence, and rate of integument evolution in early avians requires rationalization of the distribution of feather morphotypes within a phylogenetic framework (Fig. 1). To this end, statistical likelihood-based Ancestral State Estimation (ASE) methods have been applied. These methods utilize three components: a phylogenetic tree with branch lengths representing time or evolutionary rate; a set of observed character states corresponding to the tree tips (i.e. observed states in extant or fossil taxa); and an evolutionary model describing the process by which a trait changes through time. For discrete characters, this is typically a Markov model describing the transition rates between the character states. Unfortunately, ASE studies applied to early avian integument have yielded contradictory results, supporting either a single early origin of feathers around 250–240 Ma (Yang et al. 2019; Cincotta et al. 2022) or else multiple origins of filamentous integument around 160 Ma (Campione et al. 2020). Contradictory results stem from subjective methodological differences between studies relating to one of the three components of ASE.

**Figure 1.**
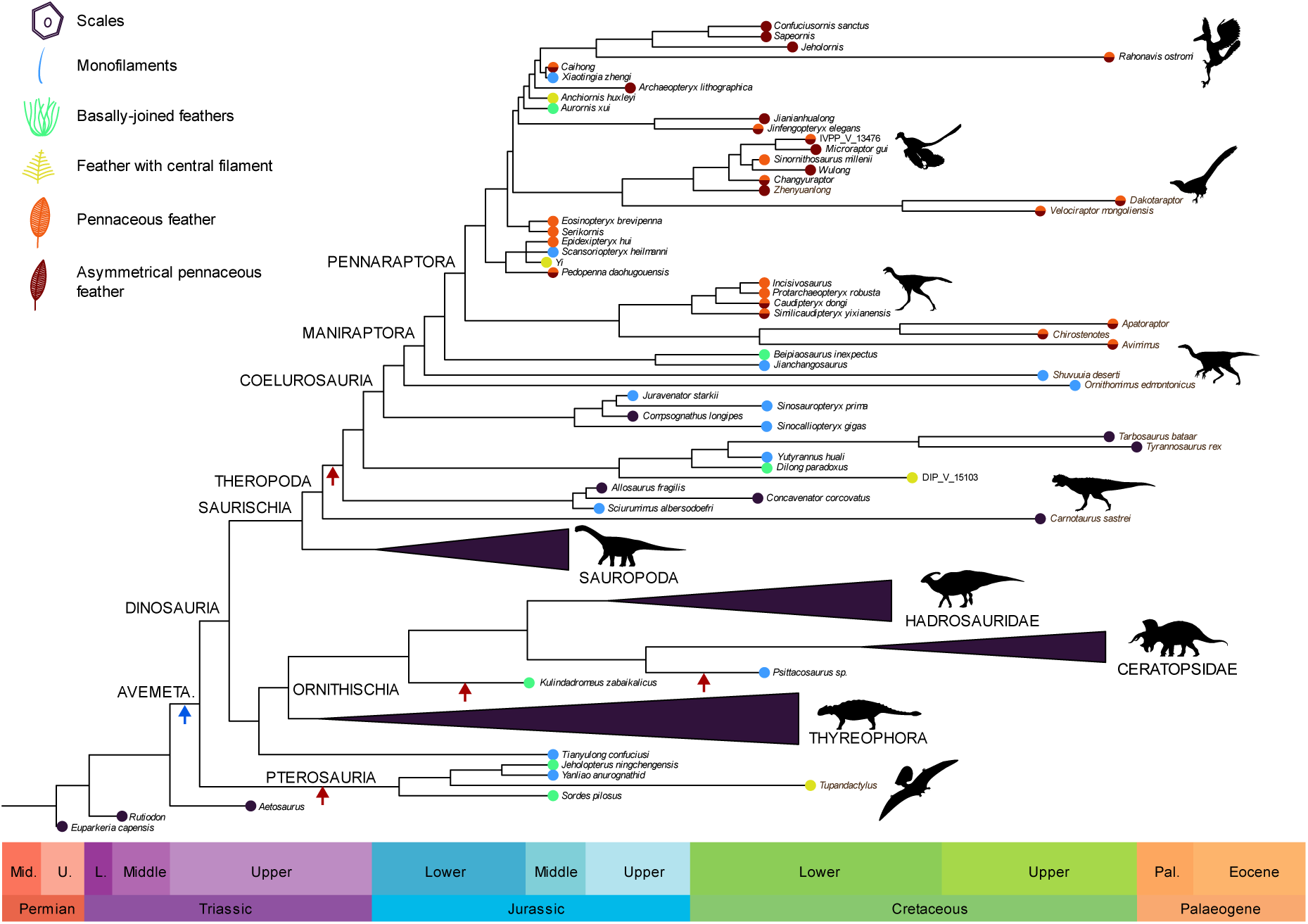
Feather evolution. The phylogenetic tree is plotted against geological time with silhouettes illustrating the main groups of dinosaurs. Drawings illustrate the feather morphotypes representative of the character states used in this study. Currently, there are two major competing hypotheses for the timing of feather origin: an early hypothesis (blue arrow) in the Triassic implying a single origin of feathers, and a late hypothesis (red arrows) in the Jurassic implying several independent origins (within Pterosauria and Dinosauria).

The first component is the phylogenetic tree. When selecting a tree, choice of outgroup(s) and branch lengths are two crucial factors to consider. Mooers and Schluter (1999) identified two major problems arising from outgroup selection: Firstly, each branch of a phylogenetic tree provides information about the rates of change. When entire clades are represented by a single lineage, as it is often the case for an outgroup, the estimates of rate may be significantly biased. Secondly, assuming transition rates are homogenous between the ingroup and the outgroup is unrealistic, particularly if the outgroup is phylogenetically distant from the ingroup. Multiple outgroups can contribute to a more robust estimation of the ancestral condition of the ingroup (Wright et al. 2015). Choice of branch length can also have a significant effect on ASE results. Previous studies, based on simulations and empirical data, have identified that ASE is sensitive to branch lengths (Litsios and Salamin 2012; Cusimano and Renner 2014). Indeed, branch lengths directly affect estimates at nodes from which they originate. Wilson et al. (2022) further confirmed these findings and suggested using model-fit statistics (such as AICc or BIC) to compare trees with alternate branch lengths set and evolutionary models. The combination associated with the best model-fit should then be selected. In ASE studies incorporating fossils, branch lengths are commonly estimated using *a posteriori* time-scaling methods such as the ‘equal’ method or the ‘minimum branch length’ (mbl) method (Laurin 2004; Brusatte et al. 2008). These methods avoid zero-length branches by either taking an equal share of preceding non-zero length branches (equal method) or by setting a minimum branch length (mbl method).

The second component is the evolutionary model. There is a plethora of evolutionary models to select from, and choosing the most appropriate is a significant challenge. The simplest model, the equal rates (ER) model, assumes that all state transitions occur at the same rate. More complex parameterized models, such as the Symmetric (SYM) and All-Rates-Different (ARD) models, relax this assumption, allowing transition rates to vary. More recently, Hidden Markov Models (HMM’s) have been applied to ASE. These models link hidden states to observed states via many-to-one mapping (i.e., multiple hidden states lead to the same observed state), permitting modelling of more complex evolutionary processes. Under the Hidden Rates Model (Beaulieu et al. 2013, Boyko and Beaulieu 2021), observed data can result from several processes occurring in different parts of the phylogeny, linked via the parameter process (Boyko and Beaulieu 2021). Structured Markov Models, proposed by Tarasov (2019, 2020), address the problem of modelling hierarchical character state dependencies, such as the classic tail color problem highlighted by Maddison (1993). Through incorporation of hidden states in a SMM, it is possible to account for both the hierarchical process resulting from character dependencies, together with the hidden process of gene regulatory evolution (Tarasov 2019). Alternatively, hierarchical characters dependencies can be modelled without hidden states by appropriate amalgamation of the rate matrices (Tarasov 2023).

The third and final component is the trait data. For discrete ASE, this comprises categorical character state observations for each tip of the tree. However, defining morphological characters and states is rarely straightforward and often subjective. Different character states can manifest in the same organism at different life stages and/or in different body regions. This is particularly true when character states are developmentally related, as is the case with feather morphotypes (e.g. Lin et al. 2020). Character states are also frequently defined with differing levels of granularity (e.g. Yang et al. 2019; Campione et al. 2020) and it is currently unclear whether it is preferable to use broad or specific character state definitions. The effect of character coding scheme on statistical ASE is poorly understood. Applying alternate coding schemes results in equivalent characters with differing numbers of states and/or different state definitions. Alternate coding schemes will therefore likely require different evolutionary models and potentially yield different evolutionary inferences.

In this study, we conduct a series of ASE experiments to evaluate the effect of outgroup sampling, time-scaling method, model selection and character coding scheme on ancestral state estimates of early avian integumentary structures. We focus on the origin(s) of feathers and the transitions between major categories of feathers defined based on their general morphologies. We benchmark the relative performance of different approaches using model fitness and a leave-one-out cross-validation approach. Through these rigorous analyses, we identify a robust set of ancestral state estimates clarifying the timing and sequence of early feather evolution. More generally, our research provides an important case-study for best practice in ASE analysis of complex traits, i.e. traits with ambiguous character states; hierarchical dependent relationships between states and/or non-linear paths of possible state transitions.

## Materials And Methods

### Data Collection and Character Coding

To conduct a phylogenetic ancestral state estimation study, we collected data on plumage characters across dinosaurs and pterosaurs from a total of 94 taxa. Character states were determined through a literature review. When more than one morphotype of feathers was reported from a taxon, only the most developmentally complex was considered. Previous analyses have shown that coding multiple trait values for taxa preserving several feather morphotypes did not allow the recovery of much information on patterns of evolution of feather types (Yang et al. 2019). This is probably because this approach does not necessarily consider similar morphotypes found in different taxa as homologous. For instance, taxa coded as having pennaceous feathers and other morphotypes were treated separately: the different combinations of feather morphotypes led to independent states. The nomenclature used for distinguishing feather types based on their general morphology follows Xu (2020). Analyses were performed using a revised version of the phylogenetic tree of Yang et al. (2019) and incorporating additional taxa by editing the tree in Mesquite (Maddison and Maddison 2023).

### Phylogenetic Time Calibration

The first appearance datum (FAD) and last appearance datum (LAD) for each taxon were determined using numerical ages for the boundaries of the stratigraphic units in which they were documented. Whenever possible, we used absolute dates reported in the scientific literature, and the International Chronostratigraphic Chart 2023-06 (Cohen et al. 2023) served as a reference otherwise. Branch lengths were then estimated using the timePaleoPhy function of the paleotree R package (Bapst 2012) and the DatePhylo function of the strap package (Bell and Lloyd 2015). The equal branch length (equal) and minimum branch length (mbl) methods were used. These equal methods (DatePhylo and timePaleoPhy) lead to distinct dated trees owing to different ordering algorithm used. In both cases, we set the root length to 1 Myr. We used the “equal_paleotree_legacy” argument in the case of timePaleoPhy. Phylogenetic trees with pie charts displaying the ancestral likelihoods were produced using the phytools R package (Revell 2012).

### Coding Schemes and Models

Ancestral states were estimated using marginal likelihood estimation. We used the corHMM function of the package of the same name (Boyko and Beaulieu 2021). First, we applied simple Markov models to each coding scheme, treating character states as independent. State transitions were modelled using the Equal Rates (ER), Symmetrical (SYM) and All Rates Different (ARD) transition rate models.

Plumage data were coded following three distinct schemes with increasing numbers of character states (Table 1). Coding #1 is formed of a simple binary character with the states “Scales” and “Feathers” (Coding #1). Coding #2 relies on a 3-state character introducing a distinction between filamentous and pennaceous feather morphotypes. Finally, Coding #3 relies on a six-state character allowing to further distinguish the main categories of feather morphology. The character matrix contains multiple character coded as uncertain in some taxa. In such instances, equal probability of the possible states was assigned.

**Table 1.** Feather coding strategies and models tested in this study. Mk = standard Markov models.

| Coding strategies | Character states | Coding schemes | State space |
| --- | --- | --- | --- |
| Coding #1 | Scales; feathers | All models: {0,1} | {0,1} |
| Coding #2 | Scales; plumulaceous feathers; pennaceous feathers | Mk: {0,1,2}<br>ED: {0,1}; {0,1}<br>SMM: {0,1}; {0,1}<br>HRM: {0,1,2} | {0,1,2}<br>{0,1,2}<br>{0&1,10,11}<br>{00,01,10,11} |
| Coding #3 | Scales; monofilamentous integument; basally joined feather; feather with central filament; pennaceous | Mk: {0,1,2,3,4,5}<br>ED: {0,1}; {0,1}, {0,1}, {0,1}, {0,1} | {0,1,2,3,4,5}<br>{0,1,2,3,4,5} |
|  | feather; asymmetrical pennaceous feather. | SMM: {0,1}, {0,1}, {0,1}, {0,1}, {0,1}<br>HRM: {0,1,2,3,4,5} | {0&1&1&1&1&1, 10000...}<br>{00000, 00001, 00010...} |

We then applied two newly developed methods for modelling dependent characters: embedded dependency (ED) and Structured Markov Models (SMMs). These two approaches require different coding schemes, and result in different state spaces. The ED model involves the amalgamation of the rate matrices of the characters (Tarasov 2023). In other words, this involves merging two or more individual characters with a dependent relationship into a single character. For instance, in the case of coding #2, the coding scheme involves a controlling character (feathers: 0, absent; 1, present) and a dependent one (feather structure: 0, plumulaceous; 1, pennaceous). Amalgamation of the rate matrices results in a single character with three states: absences of feathers (0), plumulaceous feathers (1) and pennaceous feathers (2). Two variants of the ED model can be used according to the type of character involved in the hierarchy: qualitative (ED-QL) and birth-death (ED-BD). They differ in the way the rate matrices are parameterized and whether certain transitions are permitted. The ED-QL process models the evolution of a dependent character and its inherent qualitative property. For example, using the “tail color” example of Maddison (1993), a tail may be absent or present. If the tail is present, it may be red or blue. Because tail color is an inherent property of the tail, it logically follows that when a tail evolves, it already has a color, either red or blue. This is the simplest type of embedded dependency, but another situation may arise. To illustrate the ED-BD process, Tarasov (2023) proposed the hypothetical case of tail armor evolution in species without tails, with unarmored tails and with armored tails. In this example, evolving armored tails from an ancestral state of tail absence implies two trait ‘births’: the evolution of a tail and the evolution of tail armour. Under the ED-BD model, transitions between states that would imply two or more trait ‘births’ are forbidden. As such ED-BD model assumes lags between the evolution of anatomically dependent structures.

SMMs are a type of Hidden Markov model (HMM) and assume that the evolution of the observed dependent character states is a result of an unobserved evolutionary process such as gene regulatory evolution (Tarsov 2019, 2020). SMMs are formed by amalgamating the characters as independently evolving (SMM-ind) or by imposing stricter relationships reflecting the hierarchy of the traits (SMM-switch). The approach is similar to that of the ED models, but absence states are represented by two or more hidden states. Thus, in the example of coding #2, feather absence (scales) is represented by states {00} and {01}. These hidden states can be interpreted as feather absence with a liability towards plumulaceous or pennaceous feathers respectively. The ED and SMM transition rate matrices were built using the rphenoscate package with the amaED and amaSMM functions respectively (Porto et al., 2023). Structured Markov models (SMM) were run with the argument “collapse” of corHMM set to FALSE. This setting ensures that the unobserved state combinations are modelled. Finally, we applied the hidden rates model (Boyko and Beaulieu 2021) to coding scheme 3. This model accounts for heterogeneity of the transition rates among lineages (Beaulieu et al. 2013). We used two rate categories (allowing modelling of ‘slow’ and ‘fast’ transition rates) and three rate categories (allowing modelling of ‘slow’, ‘intermediate’ and ‘fast’ rates).

Table 1 summarizes the different coding strategies and the associated coding schemes and state spaces. Diagrammatic representations of the different evolutionary models and coding schemes illustrate the states and possible transitions (Supplementary Fig. S1), as well as the associated rate parameters (Supplementary Fig. S2).

### Assessing the Sensitivity of Maximum Likelihood Analyses to a Series of Parameters

#### Experiment 1

The impact of outgroup selection was evaluated by running successive analyses with an outgroup formed of 1 fossil taxon (Outgroup #1), 2 fossil outgroups (Outgroup #2), 3 fossil taxa (Outgroup #3), and 2 fossil taxa plus a modern taxon (Outgroup #4) (Table 2). All the analyses were run with an equal tree produced with the timePaleoPhy function under unordered models. As for experiments 2 and 3 described below, the ER results do not differ strongly from the SYM results. For Experiments 1–3 we focus on SYM and ARD results in the main text, while the ER results are presented in the Supplementary Material (Tables S1–S3, Figs. S1–S3).

**Table 2.**
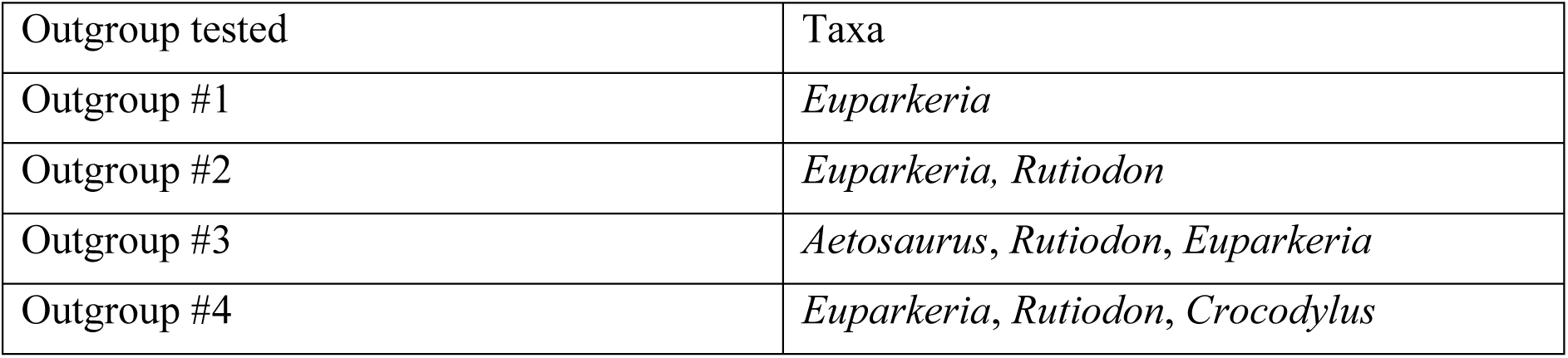
Outgroups tested in this study.

| Outgroup tested | Taxa |
| --- | --- |
| Outgroup #1 | *Euparkeria* |
| Outgroup #2 | *Euparkeria*, *Rutiodon* |
| Outgroup #3 | *Aetosaurus*, *Rutiodon*, *Euparkeria* |
| Outgroup #4 | *Euparkeria*, *Rutiodon*, *Crocodylus* |

#### Experiment 2

We tested the effect of a posteriori time-scaling methods by running the analyses with different time-scaled trees. We used the equal (DatePhylo); equal (timePaleoPhy); and mbl methods. To ensure the results are comparable, the analyses were performed with the same character (coding #3) and outgroup (outgroup #2), using the same models (unordered models).

#### Experiment 3

We also assessed the impact of the coding strategy on our results by testing three coding strategies ranging from a simple binary character to a more complex 6-state trait, as described earlier (Table 1). Analyses were run with outgroup #3 and an equal timePaleoPhy tree.

#### Experiment 4

We ran the analyses under 21 different evolutionary models, incorporating various assumptions about transition rates, rate heterogeneity and hierarchical dependent structure. As in experiment 2, all the analyses were run with the coding strategy #3 and the outgroup #3. Initially, we used an equal (timePaleoPhy) tree (see section “Effect of the Model”). We then repeated the analysis using an equal (DatePhylo) and mbl tree to ensure that our results were robust to different time scaling methods (see section “Tree/Model Combinations”).

### Measuring Model Fit, Uncertainty and Information

In all experiments, models were compared in terms of their AIC, their uncertainty and their mutual information. We used the ggplot2 R package to produce a plot comparing the models based on these statistics (Wickham 2016). Raw uncertainty (Keating 2023) is based on the raw error measure, established by Holland et al. (2020). The calculation involves summing the highest ancestral state marginal estimates for each node, dividing the result by the total number of nodes and finally, subtracting this value from 1. A theoretical absolute certainty in the ancestral state estimation would always result in a value of 0. On the other hand, the lowest possible certainty (with an equal probability of each state at each node) would be a function of the number of states: as the number of states increases, the maximal uncertainty would tend to 1. To allow comparisons between models with different state spaces, the proportion of the maximal theoretical uncertainty can be calculated. Boyko and Beaulieu (2021) defined mutual information as the difference between the unconditional entropy of the node states, and the entropy of the node states conditioned on the data. The unconditional entropy increases as the number of states in the dataset increases and defines the upper limit of what can be learned. It is set by the model and is always the same for each node of the tree. The conditional entropy is calculated using the conditional probabilities that the nodes are fixed in the various states, given the probabilities of observing the tip data. In other words, mutual information is a metric representing how much information is lost. The corHMM function can return a vector comprising the amount of information the tip states and model assign to each node. We simply summed all the values to get the total information for the whole tree.

A cross-validation approach is a more exhaustive method than AIC for computing the fitness of a model. This method involves training the model on a subset of the data and then assessing model fit with another non-overlapping subset of the observations (e.g. Lartillot, 2023). Several variants of such method exist, but we used a leave-one-out cross-validation (LOOCV) approach where each observation is removed from the dataset while the model is trained on the remaining data.

For each of the 63 model-tree combinations, we performed the following actions: (1) Turn a tip state to “?”; (2) conduct an ASE analysis; (3) Save the estimated likelihoods for the tip state; (4) Revert the tip state from “?” to its original value. The process is repeated for each of the tips of the trees, and thus results in more than 6,000 analyses in total. The argument “get.tip.states=TRUE” of the corHMM function allows us to get the estimated tip states. The LOOCV error of the model can be easily calculated for each tip by substracting the estimated likelihoods for the right tip state, from 1. This is equivalent to summing the likelihoods for all the incorrect tip states. An LOOCV mean error can then be calculated for the whole tree.

## Results

### Experiment 1: Effect of the Outgroup

The choice of outgroup does not appear to influence the ancestral state likelihoods in the case of the SYM model: the estimated ancestral likelihoods remain the same regardless of the outgroup tested (Fig. 2). The total amount of information assigned to the tree by the model + tip states remains stable, slightly above 200 bits for each of the outgroups tested (Table 3).

**Figure 2.**
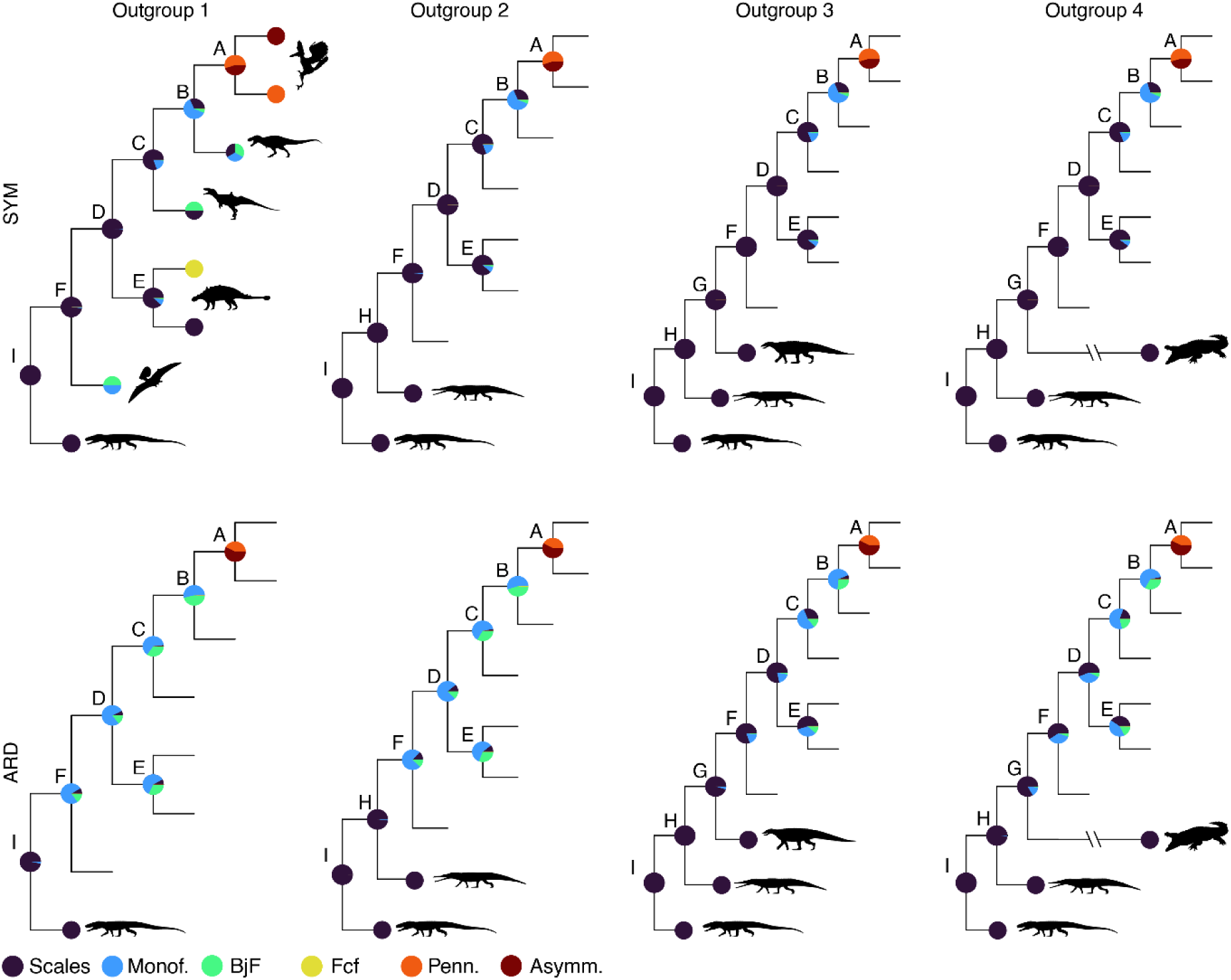
Comparison of the ancestral state likelihoods for four different outgroups selected and two transition rate models (SYM and ARD). A subset tree with only 6 nodes is plotted to facilitate the comparison. Node labels: Outgroup 1 – A= most recent common ancestor (MRCA) of *Rahonavis and Sapeornis;* B=Coelurosauria; C=Tetanurae; D=Dinosauria; E= MRCA of *Kulindadromeus* and *Psittacosaurus* sp.; F=Avemetatarsalia; G=MRCA of *Sordes pilosus* and *Aetosaurus* (Outgroup 3) or MRCA of *Sordes pilosus* and *Crocodylus* (Outgroup 4); H= MRCA of *Sordes pilosus* and *Rutiodon*; I=MRCA of *Sordes pilosus* and *Euparkeria capensis*. Abbreviations: Monof. = monofilamentous integument; BjF = basally-joined filamentous feather; FcF = Feather with central filament; Penn. = Pennaceous feather; Asymm. = asymmetrical pennaceous feather. Silhouettes from www.phylopic.org, drawn by: Oliver Demuth (*Euparkeria capensis*), Steven Traver (crocodile*, Crocodylus*), Scott Hartman (phytosaur; *Paleorhinus*), Andrew Farke (*Euoplocephalus tutus*), T. Michael Keesey (*Rahonavis ostromi*), Manuel Brea Lueiro (*T. rex*).

**Table 3.**
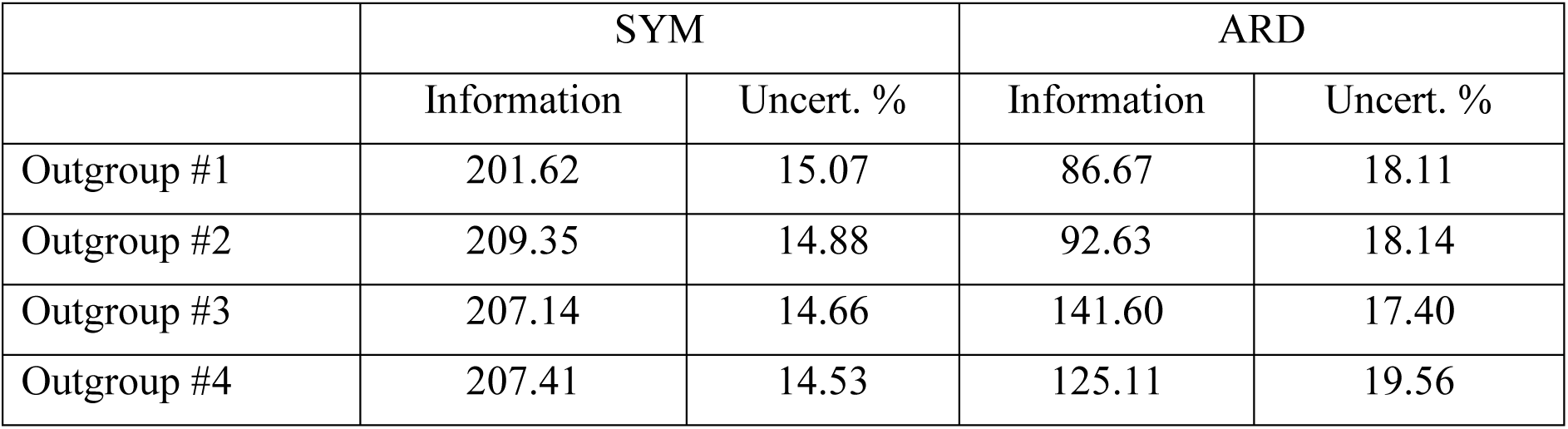
Information (in bits) and raw uncertainty calculated for each outgroup selected: with one fossil, two fossils, three fossils and, with two fossils and one extant taxon.

|  | SYM |  | ARD |  |
| --- | --- | --- | --- | --- |
|  | Information | Uncert. % | Information | Uncert. % |
| Outgroup #1 | 201.62 | 15.07 | 86.67 | 18.11 |
| Outgroup #2 | 209.35 | 14.88 | 92.63 | 18.14 |
| Outgroup #3 | 207.14 | 14.66 | 141.60 | 17.40 |
| Outgroup #4 | 207.41 | 14.53 | 125.11 | 19.56 |

The uncertainty of the results remains at about 0.12 for each tree with a different outgroup selected.

In the case of the ARD model, in contrast, outgroup choice does affect ancestral likelihoods. The result with a single outgroup was different from any of the analyses using the SYM model, noticeably placing more certainty in the presence of monofilaments early in feather evolution. Increasing the number of outgroups, either by addition of fossil taxa, or by a mix of extant and fossil taxa, changed this result, bringing the ancestral estimates closer to those proposed by the SYM models. The results suggest that ancestral estimates at nodes close to the root are particularly sensitive to outgroup selection. However, we find that this is much less the case for nodes farther from the root. For instance, the analysis recovers the same likelihoods at the node A under the ARD model, regardless of the outgroup selected (Fig. 2). At nodes D and F, the likelihoods differ strongly depending on the outgroup selected and can lead to very different conclusions regarding the absence or presence of plumage in these ancestors. Adding taxa in the outgroup seems to increase the amount of information associated with the tree under the ARD model. Using a two-taxa outgroup only leads to a limited increase of information, with respect to a tree comprising an outgroup formed of a single taxon (outgroup #2 vs outgroup #1; 93 bits and 87 bits). When a third taxon is added to the outgroup (outgroup #3) the information significantly increases to ∼142 bits. When the outgroup is formed of two fossil taxa and one extant (outgroup #4), the information decreases. The overall amount of information is much lower under the ARD model than the SYM model, irrespective of outgroup choice. The uncertainties calculated are also higher under the ARD, model with the highest uncertainty occurring for outgroup #4 (i.e. the tree with an extant outgroup).

### Experiment 2: Effect of the Time-Scaling Method

Under the equal timePaleoPhy method, internal nodes are distributed somewhat regularly along the time axis. The overall variation in the branch lengths appears lower than in the other trees. Compared to the equal timePaleoPhy method, the mbl method leads to a tree with longer internal branches, and shorter terminal branches. The equal DatePhylo method appears intermediate between the other two methods described. Some internal areas of the tree, display a high degree of similarity with the equal timePaleoPhy tree. However, there are some very short internal branches, as in the mbl method. Both the “equal datePhylo” and the “mbl” methods infer the root (avemetatarsalian node) at a younger age than the “equal tpp” method.

Our results show that time-scaling method has a large impact on the results of our ancestral state estimations (Fig. 3). Ancestral likelihoods at nodes where the model is uncertain, appear to be particularly sensitive to the time-scaling method used. Under the SYM model, there is a much higher likelihood of presence of feathers at the nodes B and C with an equal DatePhylo tree than with an equal timePaleoPhy tree.

**Figure 3.**
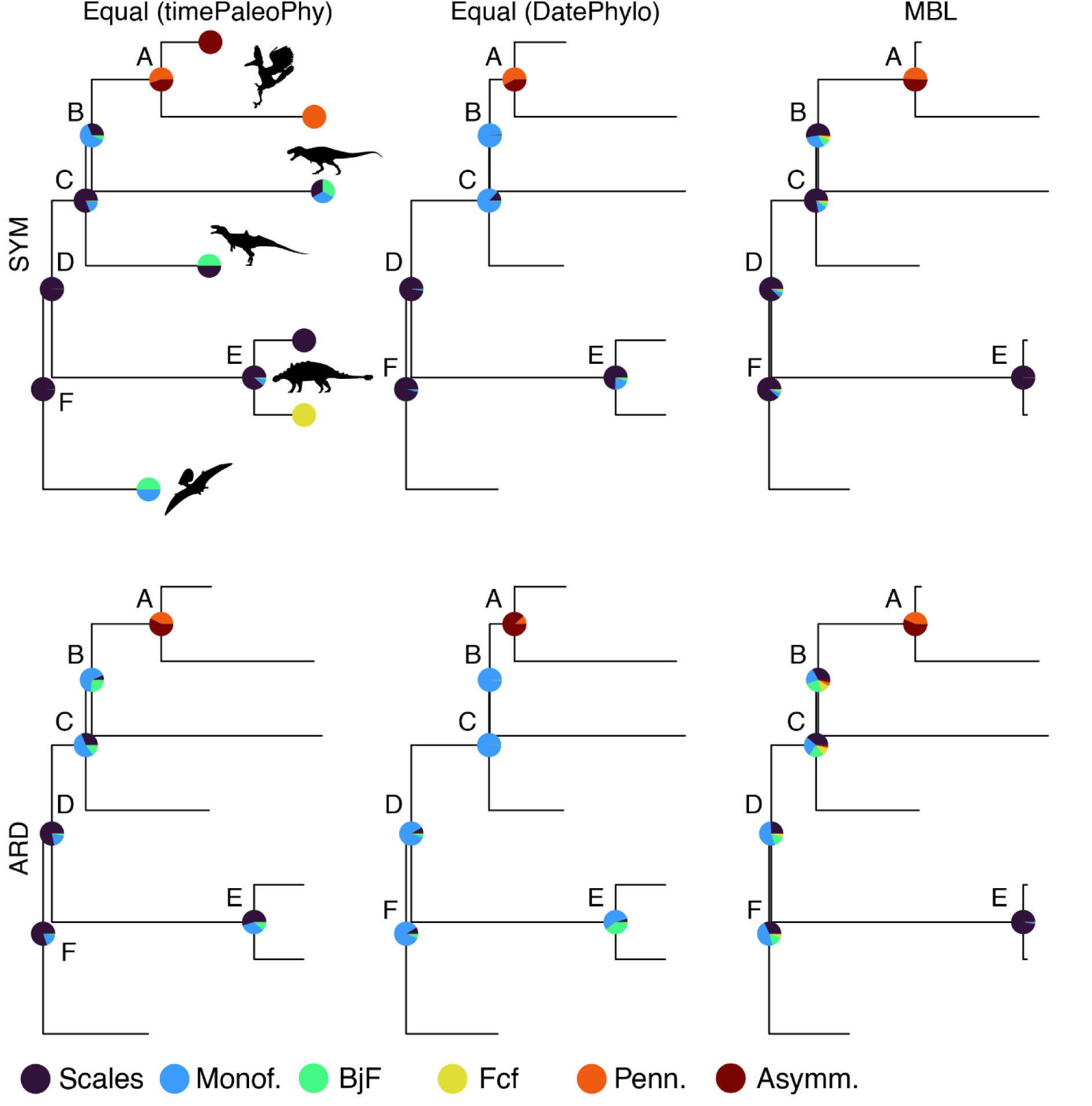
Comparison of the ancestral state likelihoods for three different trees resulting from two *a posteriori* time-scaling methods and two distinct packages (timePaleoPhy and DatePhylo). A subset tree with only six nodes is plotted to facilitate the comparison. Node labels: A=MRCA of *Rahonavis* and *Sapeornis*; B=Coelurosauria; C=Tetanurae; D=Dinosauria; E=most recent common ancestor (MRCA) of *Kulindadromeus* and *Psittacosaurus* sp.; F=Avemetatarsalia. Abbreviations in legend as defined in Fig. 2.

The total information assigned to the nodes by the model and tip data (Table 4) is maximal for the equal method under both transition rate models tested (SYM and ARD). Under SYM, both the equal timePaleoPhy and the equal DatePhylo trees, appear to have a similar total amount of information (difference of less than 4 bits). However, under ARD, the equal timePaleoPhy tree has an amount of information significantly higher (difference of about 54 bits). The uncertainty is also higher but remains significantly lower than that of the mbl tree (Table 4).

**Table 4.** Information (in bits) and uncertainty calculated for each tree obtained from different time-scaling methods. The coding #3 is used here.

|  | SYM |  |  | ARD |  |  |
| --- | --- | --- | --- | --- | --- | --- |
|  | Information | Uncert. % | AIC | Information | Uncert. % | AIC |
| Equal (DatePhylo) | 211.03 | 13.61 | 214.40 | 87.49 | 9.74 | 226.07 |
| Equal (timePaleoPhy) | 207.14 | 14.66 | 201.76 | 141.57 | 17.41 | 219.64 |
| mbl | 170.05 | 32.38 | 235.80 | 115.70 | 32.46 | 241.57 |

### Experiment 3: Effect of the Coding Strategy

The difference of AIC is the lowest for the simplest coding strategy (coding strategy 1) and maximal for the coding strategy involving the highest number of states (coding strategy 3). Coding #1 contains the least information, irrespective of the model used, while coding #3 contains the most information. It therefore seems that the number of states is positively correlated with the information content of the trait, as shown previously by Boyko and Beaulieu (2021). However, we also find a positive correlation between uncertainty of ancestral state estimates and number of character states. Raw uncertainty is the lowest for the simplest coding strategy (coding #1) and highest for the most granular coding strategy (coding #3) (Table 5).

**Table 5.** Information (in bits) assigned to the tree and raw uncertainty of the ancestral likelihoods calculated for each coding strategy tested. Abbreviations: Info.=Information (bits); Uncert.=Uncertainty.

|  | ER |  |  | SYM |  |  | ARD |  |  |
| --- | --- | --- | --- | --- | --- | --- | --- | --- | --- |
|  | AIC | Info. | Uncert. | AIC | Info. | Uncert. | AIC | Info. | Uncert. |
| **Coding #1** | 68.7 | 86.2 | 4.24 | 68.7 | 86.2 | 4.24 | 70.4 | 85.2 | 3.32 |
| **Coding #2** | 130.1 | 137.3 | 4.63 | 120.4 | 135.5 | 5.40 | 124.06 | 100.3 | 8.28 |
| **Coding #3** | 224.4 | 212.3 | 10.11 | 201.8 | 207.1 | 14.66 | 219.64 | 141.5 | 17.41 |

While all three coding strategies vary in terms of their AIC, information and uncertainty, they do not produce substantively different ancestral estimates with respect to the origin of feathers. This can be seen clearly if the likelihoods of different feather morphotypes are aggregated into a single state, equivalent to coding scheme #1. All coding schemes estimate feather absence as the most likely state at the Avemetatarsalia and Dinosauria nodes (Nodes D, F, Fig. 4). Under the SYM model, nodes B and C, farther from the root, the likelihood of feather absence seems to increase as the coding strategy becomes more granular. A somewhat opposite pattern is observed under ARD. At the node E, increasing the number of states increases the likelihood of presence of plumage.

**Figure 4.**
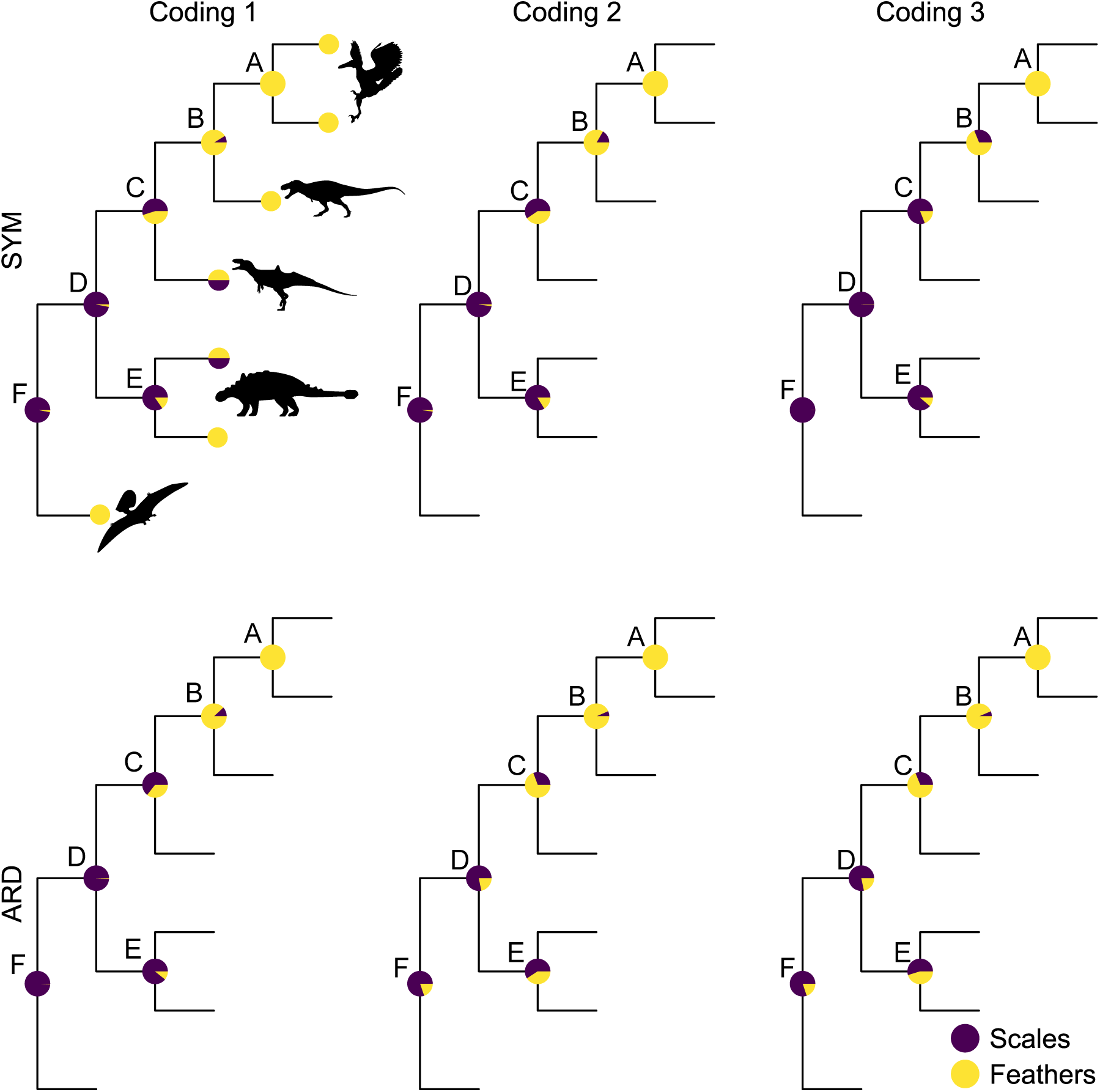
Comparison of the ancestral state likelihoods obtained for three different coding strategies. A subset tree with only 6 nodes is plotted to facilitate the comparison. For each coding strategy, the likelihoods of absence and presence of plumage are displayed through the aggregation of the likelihoods of the five feather morphotype states. This is effectively reducing coding strategies #2 and #3 to coding strategy #1, so that all the results can be compared. Node labels: A= most recent common ancestor (MRCA) of *Rahonavis* and *Sapeornis*;. B=Coelurosauria; C=Tetanurae; D=MRCA of *Kulindadromeus* and *Psittacosaurus* sp.; E=Dinosauria; F=Avemetatarsalia.

### Experiment 4: Effect of the Model

We estimated ancestral states under three transition models (ER, SYM and ARD) and seven different Markov model architectures (Unordered, Ordered, ED, SMM-ind, SMM-sw, HRM(2) and HRM(3)) resulting in 21 models (Supplementary Table S3). The results of these analyses are summarized using a simplified version of the phylogenetic tree in Fig. 5.

**Figure 5.**
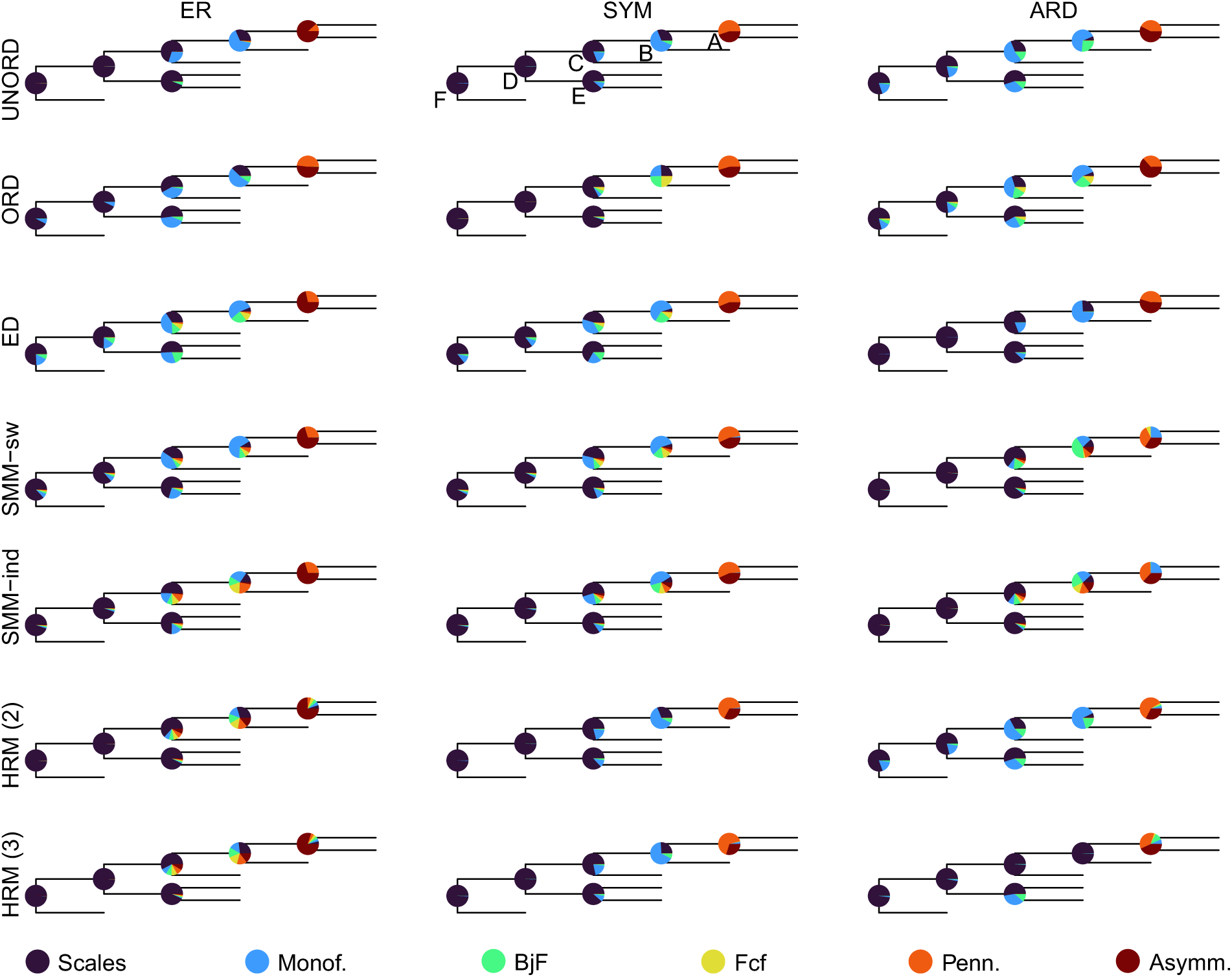
Comparison of the ancestral state likelihoods for seven different models under ER, SYM and ARD. A subset tree with only six nodes is plotted to facilitate the comparison. Node labels: A= most recent common ancestor (MRCA) of *Rahonavis* and *Sapeornis*;. B=Coelurosauria; C=Tetanurae; D=MRCA of *Kulindadromeus* and *Psittacosaurus* sp.; E=Dinosauria; F=Avemetatarsalia. Abbreviations: ED = embedded dependency; SMM-sw = structured Markov model, switch; SMM-ind = structured Markov model, independent; HRM (2) = Hidden rates model with 2 rate categories; HRM (3) = Hidden rates model with 3 rate categories. Abbreviations as in Figures 2 and 3.

All the 21 models tested support the hypothesis of several independent origins of feathers throughout their evolutionary history. The presence of scales in the avemetatarsalian ancestor of dinosaurs and pterosaurs is inferred with likelihoods ranging from about 76% (ER ED) to more than 99% (ER UNORD). Most of the models also infer the absence of feathers at node C, although a few suggest their possible presence (e.g. ER ORD, ER ED, ARD UNORD). The hidden Markov models (SMM and HRM) tend to be uncertain regarding the ancestral state at node B, and to some extant at node C, as well.

Using AIC for model selection, the ARD ORD model is best at explaining the data observed. However, the model returning results with the lowest uncertainty is ER UNORD. The SMM-sw model is associated with a higher uncertainty but with a very high information. For the standard Markov models (UNORD and ORD models), the amount of information is nearly the same when the ER or SYM transition rate models are used but decreases significantly under the ARD model. In addition, ordered models show a lower information content of the tree than unordered models. This is particularly marked with the ARD models, where the information decreases by more than 26 bits. Ordered models are also associated with a higher uncertainty of the results. Here, the effect is strongest under an ER model where the uncertainty nearly doubles with ordering. Among all the 21 models tested the lowest uncertainty of results is achieved with the ER unordered model, which is the simplest model tested with only a single parameter to estimate. Its AIC is among the highest (indicating a poor fit of the model to the data) observed in this study.

In the case of the embedded dependency models, the information assigned to the tree is maximal under ARD. Under ER and SYM, on the other hand, the ED models are associated with the lowest amounts of information (i.e. nearly half that of the other models tested). The SMM models are associated with very high uncertainties of the results under ARD. This uncertainty is much lower under the ER and SYM models, while the information content of the trees is high.

In the case of the HRM models, the amount of information is nearly the same under ER and SYM, and significantly lower under ARD. Here, the uncertainty does not seem to be correlated with the number of rate categories, or even the transition rate model used. In fact, the uncertainty is even higher under ER than SYM or ARD which is not what is observed with the other models. The HRMs also do not seem to extract more information than the other Markov models.

### Tree/Model Combinations

In the previous section we tested the models using the equal timePaleoPhy tree. However, different trees may be favored under different models. To test whether this is the case here, we ran additional analyses with the equal DatePhylo and mbl trees, thus resulting in 63 tree/model combinations (Supplementary Table S4). Eight combinations inferred the presence of plumage at the avemetatarsalian node. These are all associated with low information, and except for two combinations, have a moderate to high uncertainty of the ancestral state estimates. The favored model is function of the metrics used: AIC (ARD ORD, mbl tree), or LOOCV mean error (ARD HRM (3), equal (DatePhylo) tree).We calculated Akaike weights (Akaike, 1978) and followed a similar approach for the LOOCV mean error (see Supplementary text). The ancestral likelihoods resulting from this model averaging approach, whether they rely exclusively on Akaike weights (Supplementary Fig. S6), or LOOCV mean error weights, do not lead to different evolutionary interpretations.

### Cross-validation approach

The LOOCV results (Supplementary Table S5) show that the mean error of the models ranges from approximately 0.32 to 0.46. Some models associated with high AIC values (e.g. ARD HRM 3 with an equal DatePhylo tree; 324.7), have in fact the lowest LOOCV mean errors out of the models tested (around 0.32). All the ARD HRM models are thus slightly outperforming or nearly equal to the ARD ORD models although they have much higher AIC values. There is little correlation between AIC and LOOCV mean error (Fig. 6). Furthermore, information and uncertainty do not seem to be correlated with LOOCV mean error. Some models with low information have either low mean errors (ARD HRM models with an equal DatePhylo tree) or high mean errors (e.g. ARD SMM models with an equal DatePhylo tree). A similar observation can be made with the uncertainty metric. Models associated with low uncertainties, such as the ER UNORD model, also have high LOOCV mean errors.

**Figure 6.**
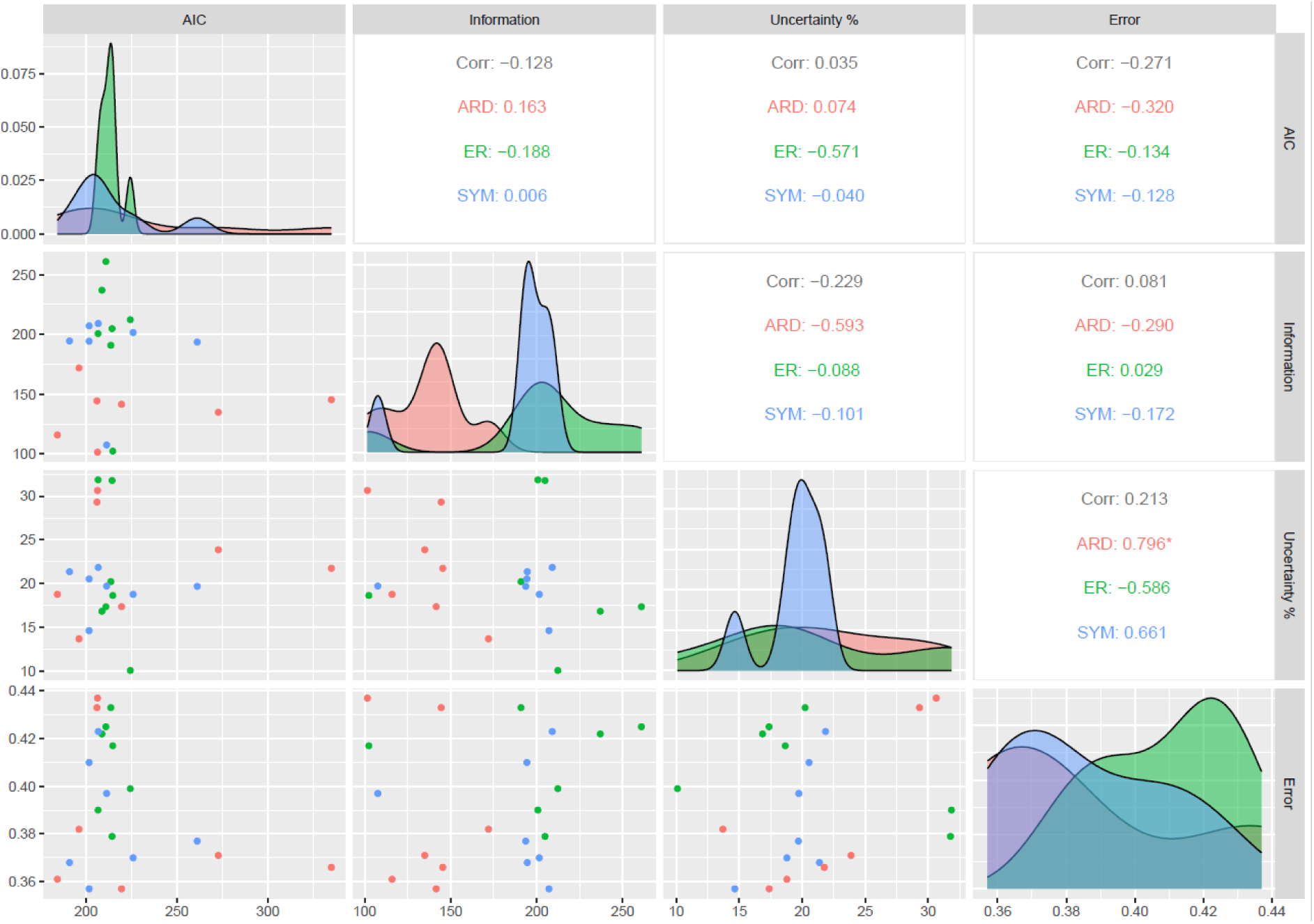
Comparison of the 21 models tested in this study using the AIC, Information, Uncertainty and LOOCV mean error of the models.

## Discussion

### Outgroup Importance

Our results show that choice of outgroup has little influence on ancestral state estimates under the SYM model. The ancestral likelihoods appear identical across the analyses performed with three different outgroups, but this is not the case under the ARD model. The analyses performed show that increasing the number of taxa comprising the outgroup changes the ancestral likelihoods at key nodes. While the analysis performed with an outgroup comprising a single taxon or two taxa, estimate the presence of feathers at the root of the tree (i.e., avemetatarsalian node), analyses with an outgroup comprising three taxa estimate a higher likelihood of absence of any plumage at that node. This implies that, in some cases, interpretations surrounding the evolution of a trait can depend on the outgroup selected. For instance, Yang et al. (2019) and Cincotta et al. (2022) proposed the hypothesis of a single point of origin of feathers around 240 Ma at the avemetatarsalian node, based on maximum likelihood methods. In both studies, an ARD model was selected, and an outgroup formed of a single taxon (Yang et al., 2019) was used, or the analysis was conducted without an outgroup (Cincotta et al., 2022). Including several outgroups may allow estimation of the ancestral states with a higher confidence at some nodes close to the root. However, adding numerous outgroups may also involve a risk of biasing the rate estimation. Such issues are of particular importance in the case of ASE analyses of feather evolution. As mentioned earlier, previous studies have led to contradictory results regarding the origin(s) of feathers. While some analyses recovered the presence of filamentous integument in the ancestor of pterosaurs and dinosaurs (Avemetatarsalia), others found several independent origins within Pterosauria and Dinosauria. The avemetatarsalian node is close to the root of the phylogenetic tree and therefore, ancestral state estimates at this node are potentially very sensitive to outgroup selection. We suggest selecting multiple outgroups based on phylogenetic proximity to the ingroup and assessing the effect of outgroup selection on the analyses by exploring how the ancestral estimates of basal nodes vary with alternative outgroups.

### Time-Scaling Importance

Several studies have shown that ASE is sensitive to branch lengths and thus, to the a posteriori time-scaling method used (e.g. Bapst 2014; Bapst and Hopkins 2017). Our results support this assertion. The information and the uncertainty vary according to the method used. The time-scaling methods impact the position of the internal nodes along the x-axis and the branch lengths. The same model can lead to different results when various trees are used (see Supplementary Table S4). The equal timePaleoPhy tree is characterized by internal nodes more evenly distributed along time than the other two methods tested. In the equal DatePhylo tree, some branches are elongated or shortened resulting in an apparent shift toward the tips of a series of internal nodes. This is also the case for the mbl tree, but even more pronounced. Within Ornithischia some internal branches and long and multiple nodes are placed much closer to the tips than in the other trees. We recommend testing different combinations of models and time-scaling methods. Leave-one-out cross-validation can then be used to select the best combination. Prior knowledge of the biological processes studied may also guide the selection of the best tree and model combination. Model averaging can be used to incorporate the uncertainty from different time-scaling methods.

### Coding Strategy

All the coding strategies tested retained sufficient historical information on the trait studied. Logically, increasing the number of states increased the information assigned by the model and tip data to the tree nodes (Table 5). This is also positively correlated with the uncertainty of the results. Under both SYM and ARD, the coding strategy seems to affect the estimated likelihoods of presence of plumage at some nodes (Fig. 3). However, the impact remains limited, and the overall interpretation of the evolutionary history did not change. These results suggest that coding strategies are less of a concern than we had feared; the definitions of states and their granularity do not necessarily lead to different macroevolutionary inferences.

### Model Selection

AIC scores and LOOCV mean error scores can both be used to estimate the relative fitness of a model. However, our results show little correlation between these measures, raising questions over their reliability. As such, it is worth considering what these scores are actually measuring: AIC uses a model’s likelihood estimation as a measure of fit, which can be inflated by model overfitting, particularly for small sample sizes. AIC employs parsimony to penalize more complex models in proportion to the number of parameters, yet this may not be sufficient to overcome overfitting. In contrast, LOOCV explicitly measures how well the model generalizes over unseen data, which inherently provides a more robust assessment against overfitting (Lartillot 2023). As such, we are inclined to favor the LOOCV mean error scores over AIC.

The LOOCV results suggest that the best fitting models are the SYM and ARD models (ordered and unordered), as well as the SYM and ARD HRM models. The SMM are associated with the highest LOOCV mean errors: between approximately 0.42 and 0.45. SMM involve much more free parameters than standard Markov models and as such, are strongly data dependent (Tarasov 2019). Beaulieu et al. (2013) found that trees with a minimum of 60–120 taxa are required so that SMM can be used in ASE. However, the ER models (unordered and ordered) are similarly associated with high LOOCV mean errors (about 0.40 and 0.43). This implies that these models are strongly biased: with a single parameter, they are too simple to accurately model feather evolution and explain the data observed.

### Implications for Feather Evolution

Most of the models tested support the hypothesis of feathers originating several times throughout their evolutionary history. Only eight model-tree combinations out of 63 tested infer the presence of feathers at the avemetatarsalian node with a likelihood above 0.4. These are associated with low information. The ARD ordered models selected based on the AIC, suggest a minimum of three independent points of origin within Pterosauria, Saurischia and Ornithischia. A similar observation can be made with an average of the models using weightings based on the LOOCV mean error of the models (Fig. 7). If that is the case, this would correspond to a late origin with filaments homologous to modern bird feathers evolving between 190 and 212 million years ago, as opposed to an early origin of feathers suggested by the results of previous ASE analyses (Yang et al. 2019; Cincotta et al. 2022).

**Figure 7.**
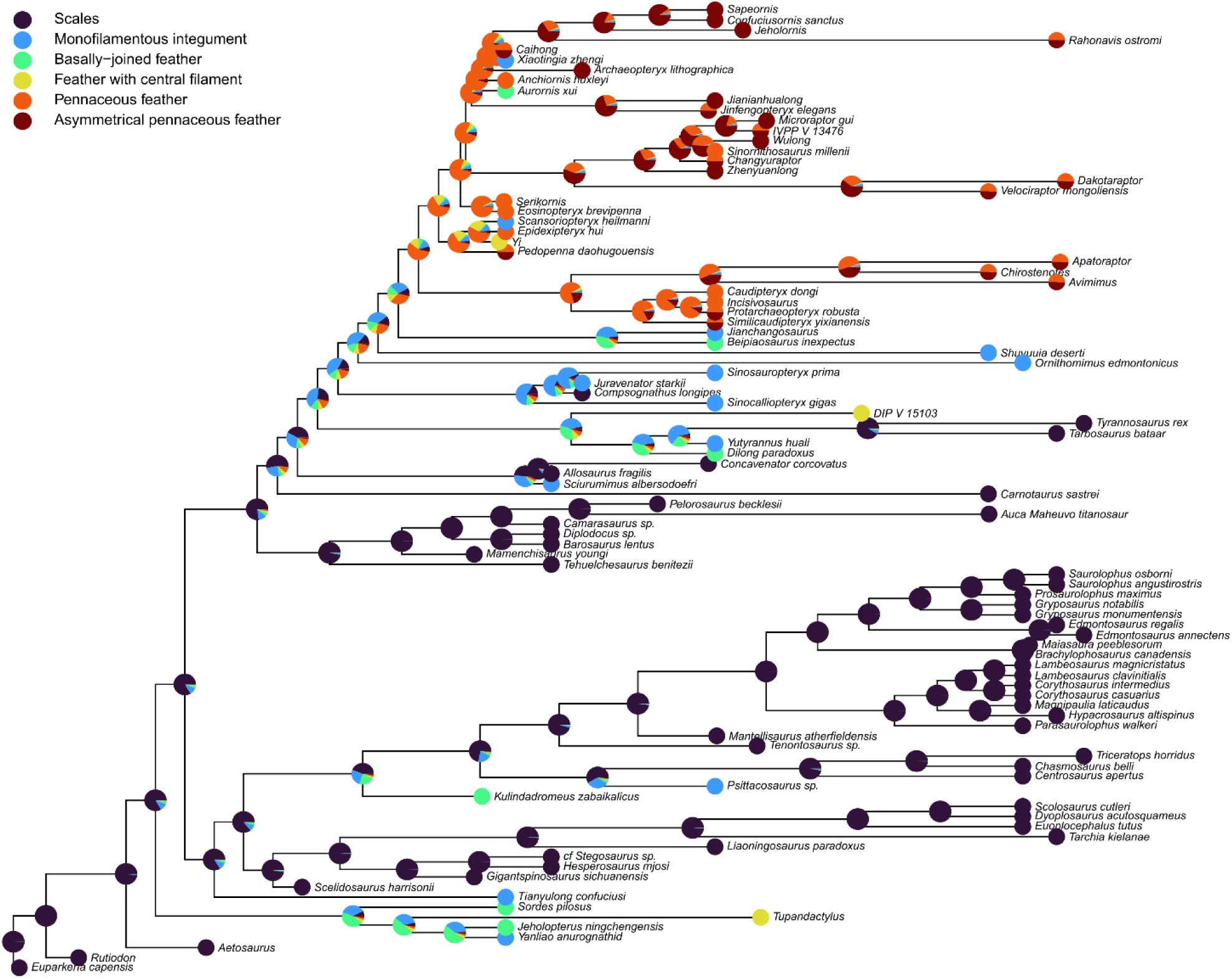
Average of the 63 model/tree combinations using weightings based on the error of the models. The ancestral likelihoods are plotted on the equal (timePaleoPhy) tree.

Pennaceous feather morphotypes have been reported on the forelimb and tail, from basal and more derived paravian taxa, and from Oviraptorosauria. Our analyses support the hypothesis that pennaceous feathers were inherited from a common ancestor of Paraves and Oviraptorosauria, in the Lower Jurassic (∼185 Ma). Based on the presence of quill-knobs, pennaceous feathers have been inferred in a specimen of *Ornithomimus* (Zelenitsky et al. 2012). However, the preserved morphology would rather suggest that these are broad monofilamentous feathers (Xu 2020). Feather specialization with short coverts overlapping longer flight feathers is observed in Dromaeosauridae (e.g., *Microraptor, Changyuraptor*, *Zhenyuanlong*) and in basal avian taxa (*Archaeopteryx*, *Jeholornis*, *Sapeornis*, *Confuciusornis*). The asymmetry of flight feathers follows a similar evolutionary pattern: asymmetrical feathers were observed in dromaeosaurid dinosaurs (e.g. *Zhenyuanlong, Wulong*, *Microraptor*) and Avian taxa. Since the forelimb plumage is not preserved in troondontid specimens, it is unclear whether feather specialization and asymmetry occurred separately in Dromaeosauridae and Aves (i.e. convergent evolution) or was inherited from a common ancestor.

Although these results seem to refute the hypothesis of feathers being a synapomorphy of Pterosauria and Dinosauria, further investigation is required to clarify the early evolutionary history of feathers. Some of the models tested in this study, in particular the Hidden Markov models, may perform better with a tree comprising additional tips. Furthermore, a major limitation of recent studies including this one, is related to the way the plumage information is coded. Only the most developmentally complex morphotype is considered even when several are observed within a taxon. A pre-requisite for significantly improving future ASE analyses may involve developing new ways to include such information. A possible solution requires using models in which polymorphic states are included and treated as intermediate between the corresponding monomorphic conditions.

## Conclusions

Feather evolution served as an ideal case study to explore different statistical approaches in ancestral state estimation of complex characters. We tested the effect of different outgroups, branch lengths, coding strategies, and models on our results. All these parameters influenced the end results on their own or in association. Among the models tested, ARD ORD is associated with the lowest AIC, theoretically suggesting the best model fit. However, a cross-validation approach revealed that some models with a higher AIC are associated with similar or even slightly lower errors. This would suggest that model selection using AIC might not provide a sufficient approximation of the absolute fit when the sample size (number of tips) is limited. Model averaging using different AIC or LOOCV mean error weightings provides similar overall ancestral state estimates. The results support the hypothesis of several independent origins of feathers.

Ultimately these results inform best practices to adopt when modelling complex discrete characters. It is critical to carefully establish the appropriate coding strategy allowing to provide the maximum information on the processes studied. The outgroup selection and branch lengths must be carefully considered as well. Finally, analyses should be conducted with several statistical methods. The AIC may not be appropriate for model selection for small datasets, which are typical for palaeontological studies: it is preferrable to compare the models using cross validation.

## Acknowledgments

We thank Sergei Tarasov and James Boyko for kindly providing advice on conducting the analyses.

