## Supplementary Materials for "Estimating Ancestral States of Complex Characters: A Case Study on the Evolution of Feathers"

**Supplementary text: model averaging.** We calculated Akaike weights (Akaike, 1978). The first step involves, for each tree-model combination, the calculation of the differences in AIC, with the best combination.

In the case of AIC and uncertainty, this can be written:

$$\Delta_i(AIC) = AIC_i - \min AIC$$

The Akaike weights can be calculated as:

$$w_i(AIC) = \frac{\exp(-0.5\Delta(\mathrm{AIC}))}{\sum_{k=1}^{K} \exp(-0.5\Delta(\mathrm{AIC}))}$$

The ancestral likelihoods of each tree/model combination are then multiplied by the corresponding weighting and summed across the 63 combinations. This results in averaged ancestral likelihoods. We calculated the weightings and plotted the ancestral likelihoods for AIC (Supplementary Figure S2).

In the case of the Error, the weights were calculated by subtracting the error of a model from 1 and dividing the result by the sum of the errors of all the models.

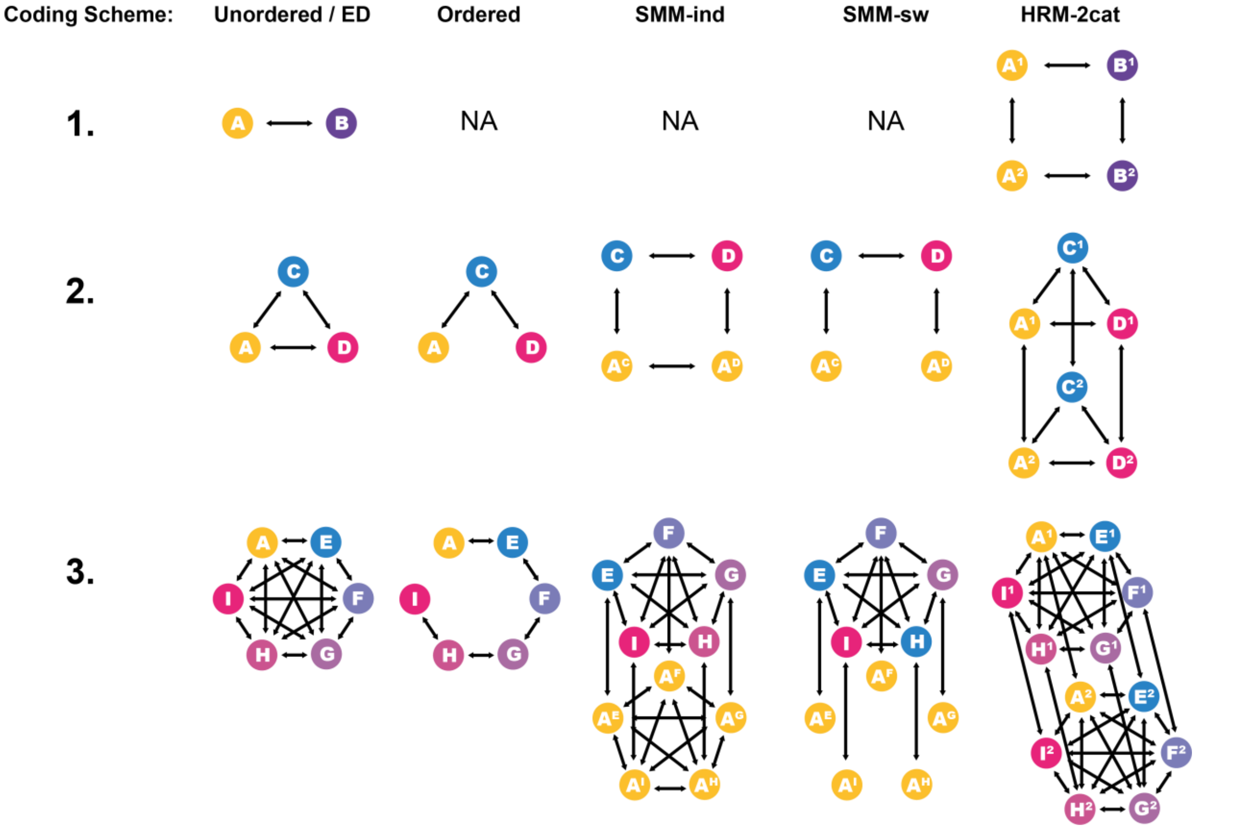

FIGURE S1. Diagrammatic representations of the different evolutionary models and coding schemes. Each letter refers to a state while the arrows show the possible transitions. A=Scales; B=Feather; C=filamentous feather; D=Pennaceous feather; E=Monofilamentous integument; F=Basally-joined feather; G=feather with central filament; H=Pennaceous feather; I=Asymmetrical pennaceous feather. Superscript letters are used in the hidden layer of the SMM models to denote liabilities. Superscript numbers in the HRM model indicate the rate categories (either 1=slow or 2=fast). For clarity, the HRM-3cat model is not shown on this figure.

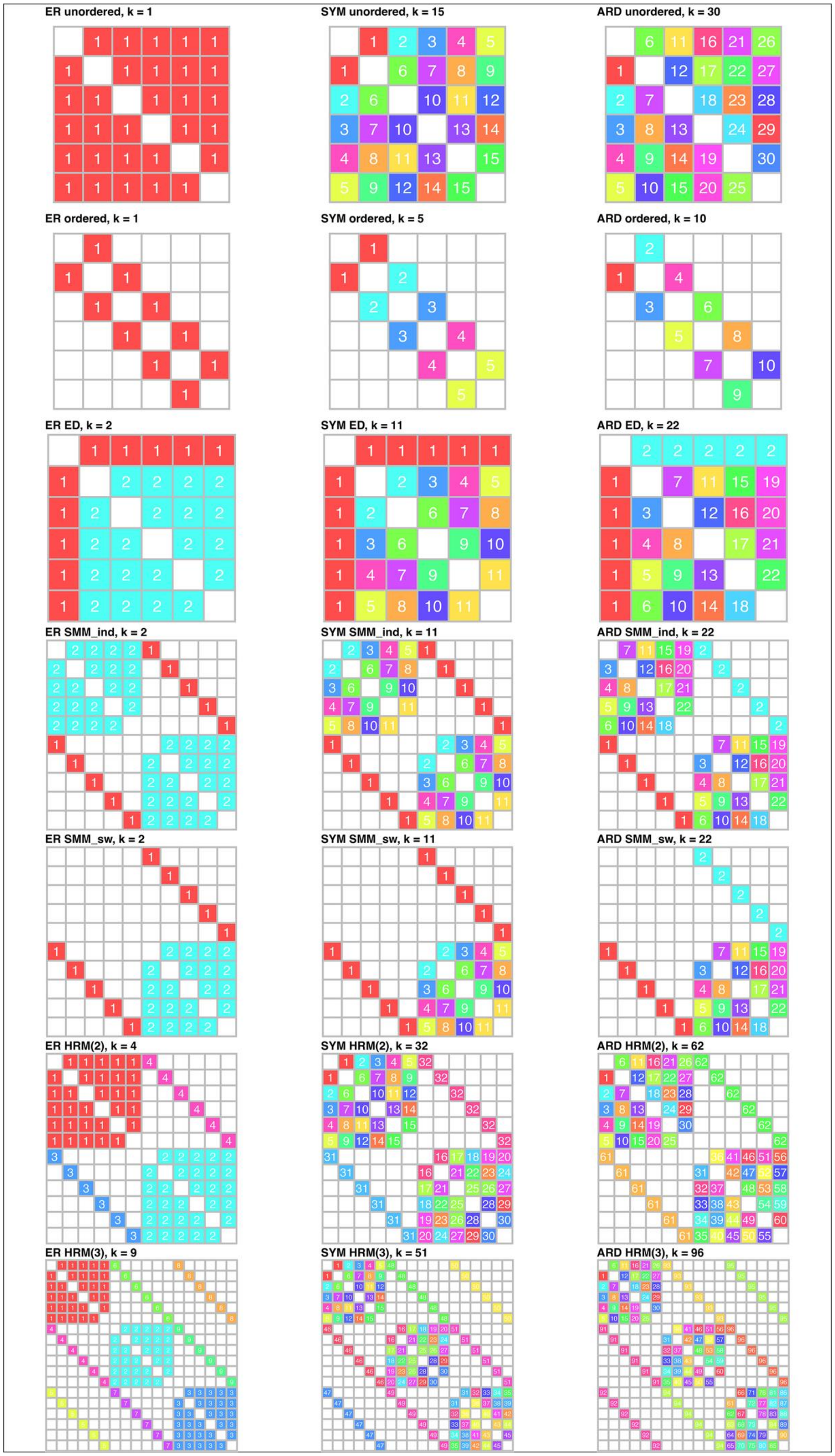

FIGURE S2. Diagrammatic representations of the 21 Markov Models tested in Experiment 4. k = number of rate parameters.

| | ER | | SYM | | ARD | |
|---|---|---|---|---|---|---|
| | Info. | Uncert. % | Info. | Uncert. % | Info. | Uncert. % |
| Outgroup #1 | 206.59 | 10.46 | 201.62 | 15.07 | 86.67 | 18.11 |
| Outgroup #2 | 201.62 | 10.30 | 209.35 | 14.88 | 92.63 | 18.14 |
| Outgroup #3 | 212.27 | 10.11 | 207.14 | 14.66 | 141.60 | 17.40 |
| Outgroup #4 | 213.56 | 09.71 | 207.41 | 14.53 | 125.11 | 19.56 |

TABLE S1. Outgroup effect.

| | ER | | | SYM | | | ARD | | |
|---|---|---|---|---|---|---|---|---|---|
| | Info | Uncert % | AIC | Info. | Uncert. % | AIC | Info. | Uncert. % | AIC |
| Equal (DatePhylo) | 214.84 | 8.74 | 240.31 | 211.03 | 13.61 | 214.40 | 87.49 | 9.74 | 226.07 |
| Equal (timePaleoPhy) | 212.27 | 10.11 | 224.4 | 207.14 | 14.66 | 201.76 | 141.57 | 17.41 | 219.64 |
| mbl | 185.94 | 20.21 | 260.74 | 170.05 | 32.38 | 235.80 | 115.70 | 32.46 | 241.57 |

TABLE S2. Time-scaling effect.

| | **ER** | | | **SYM** | | | **ARD** | | |
|---|---|---|---|---|---|---|---|---|---|
| | Uncert. | Info. | AIC | Uncert. | Info. | AIC | Uncert. | Info. | AIC |
| **UNORD** | 0.084 | 212.27 | 224.4 | 0.122 | 207.14 | 201.8 | 0.145 | 141.60 | 219.6 |
| **ORD** | 0.169 | 190.90 | 213.7 | 0.178 | 194.52 | 191.0 | 0.157 | 115.99 | 184.3 |
| **ED** | 0.155 | 102.49 | 214.8 | 0.164 | 107.70 | 211.4 | 0.114 | 172.00 | 196.2 |
| **SMM-sw** | 0.145 | 260.88 | 211.0 | 0.182 | 209.06 | 206.8 | 0.255 | 101.62 | 206.4 |
| **SMM-**[illegible] | 0.140 | 236.97 | 208.8 | 0.171 | 194.31 | 201.8 | 0.244 | 144.49 | 206.1 |
| **HRM (2)** | 0.265 | 200.70 | 206.7 | 0.157 | 201.55 | 226.0 | 0.199 | 134.93 | 272.8 |
| **HRM (3)** | 0.265 | 204.85 | 214.4 | 0.164 | 193.65 | 261.2 | 0.181 | 145.45 | 335.0 |

TABLE S3. Information (in bits), uncertainty and AIC calculated for each model tested in Experiment 4. Abbreviation: Info.= Information (bits); Uncert.=Uncertainty.

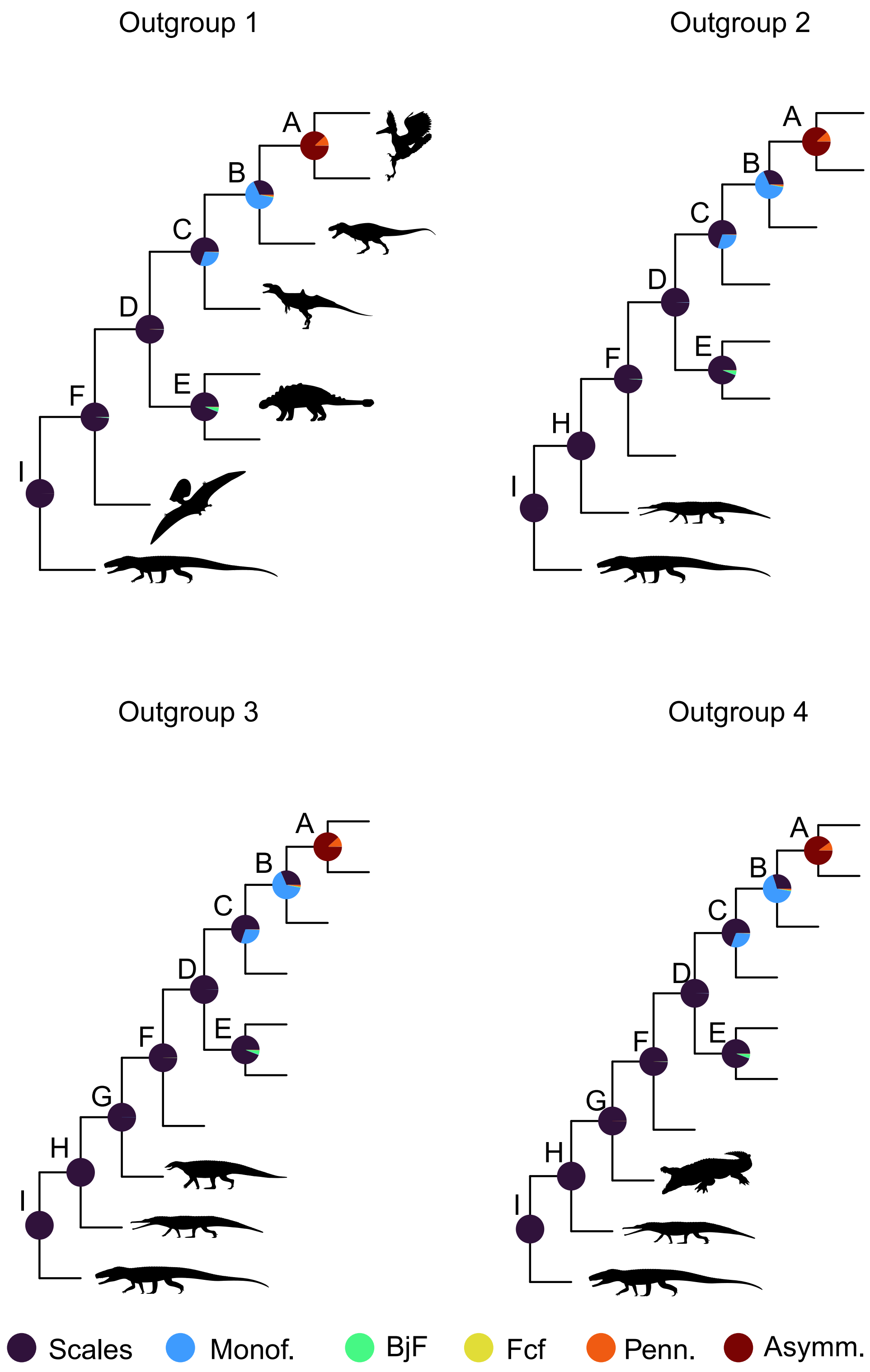

FIGURE S3. Comparison of the ancestral state likelihoods under an ER transition rate model, for four different outgroups.

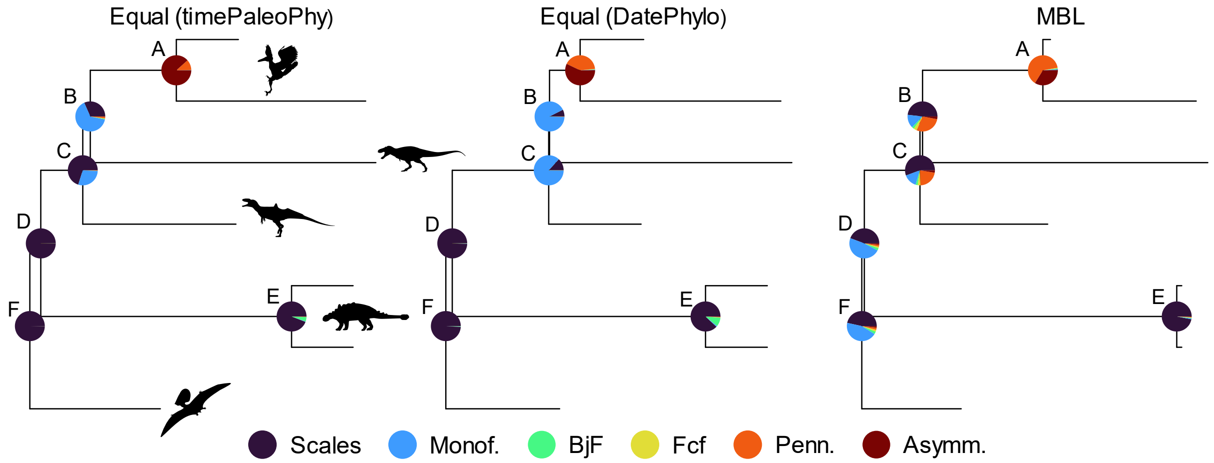

FIGURE S4. Comparison of the ancestral state likelihoods under an ER transition rate model, for three different trees resulting from two a posteriori time-scaling methods and two distinct packages (timePaleoPhy and DatePhylo).

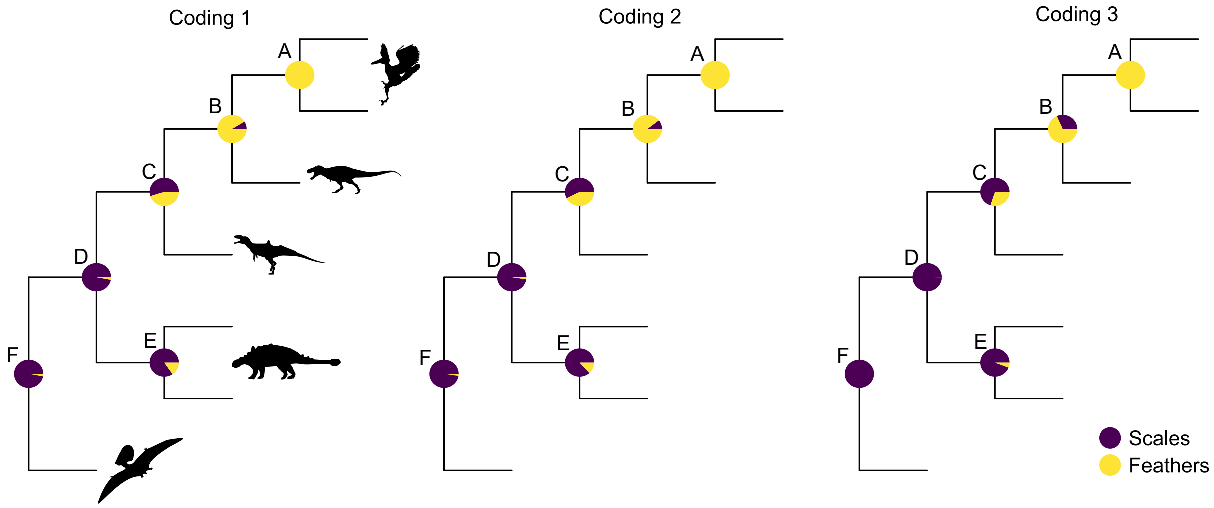

FIGURE S5. Comparison of the ancestral state likelihoods under an ER transition rate model, for three different coding strategies.

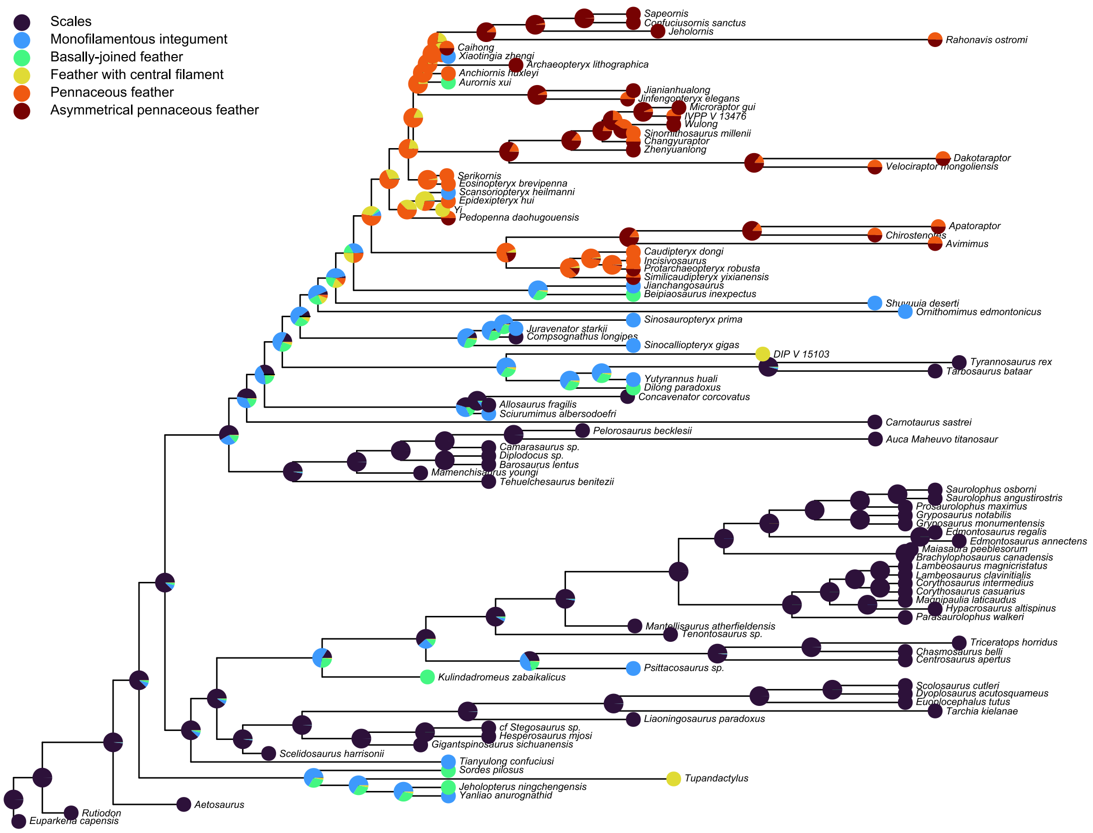

FIGURE S6. Average of the 63 model/tree combinations using weightings based on AIC. The ancestral likelihoods are plotted on the equal (timePaleoPhy) tree.

| | ER | | | | SYM | | | | ARD | | | |
|---|---|---|---|---|---|---|---|---|---|---|---|---|
| | Information | Uncert % | AIC | Avem. | Information | Uncert % | AIC | Avem. | Information | Uncert % | AIC | Avem. |
| **UNORD** | | | | | | | | | | | | |
| Equal tpp | 212.27 | **10.11** | 224.4 | 0.997 | 207.14 | 14.66 | 201.8 | 0.995 | 141.60 | 17.39 | 219.6 | 0.803 |
| Equal dp | 214.84 | **8.74** | 240.3 | 0.991 | 211.03 | 13.61 | 214.4 | 0.962 | 87.49 | 9.74 | 226.1 | 0.087 |
| mbl | 211.55 | **10.49** | 227.9 | 0.992 | 207.19 | 14.83 | 201.5 | 0.993 | 152.85 | 18.76 | 218.8 | 0.930 |
| **ORD** | | | | | | | | | | | | |
| Equal tpp | 190.9 | 20.24 | 213.7 | 0.925 | 194.52 | 21.38 | 191.0 | 0.998 | 115.99 | 18.79 | 184.3 | 0.790 |
| Equal dp | 194.17 | 17.85 | 245.3 | 0.870 | 188.84 | 25.40 | 206.2 | 0.990 | 88.09 | 20.95 | 188.5 | 0.054 |
| mbl | 190.46 | 21.70 | 205.1 | 0.907 | 190.40 | 18.82 | 190.3 | 0.995 | 146.40 | 19.97 | 181.6 | 0.926 |
| **ED** | | | | | | | | | | | | |
| Equal tpp | 102.49 | 18.66 | 214.8 | 0.758 | 107.70 | 19.74 | 211.4 | 0.858 | 172.00 | 13.72 | 196.2 | 0.997 |
| Equal dp | 101.61 | 18.91 | 234.3 | 0.228 | 98.60 | 24.02 | 228.4 | 0.336 | 147.12 | 13.53 | 211.2 | 0.952 |
| mbl | 109.40 | 16.20 | 220.5 | 0.356 | 105.75 | 19.49 | 220.7 | 0.336 | 169.25 | 17.39 | 200.1 | 0.995 |
| **SMM-sw** | | | | | | | | | | | | |
| Equal tpp | **260.88** | 17.38 | 211.0 | 0.871 | 209.06 | 21.86 | 206.8 | 0.925 | 101.62 | 30.64 | 206.4 | 0.989 |
| Equal dp | **264.44** | 20.41 | 227.6 | 0.656 | 197.76 | 25.64 | 217.7 | 0.813 | 95.20 | 37.28 | 205.3 | 0.830 |
| mbl | **256.79** | 16.15 | 214.5 | 0.818 | 205.13 | 24.65 | 212.5 | 0.824 | 107.28 | 32.22 | 203.5 | 0.950 |
| **SMM-ind** | | | | | | | | | | | | |
| Equal tpp | 236.97 | 16.86 | 208.8 | 0.950 | 194.31 | 20.55 | 201.8 | 0.970 | 144.49 | 29.31 | 206.1 | 0.989 |
| Equal dp | 254.36 | 17.03 | 225.2 | 0.878 | 212.35 | 22.48 | 216.0 | 0.882 | 98.67 | 36.95 | 205.3 | 0.830 |
| mbl | 243.42 | 14.11 | 212.3 | 0.927 | 206.11 | 21.05 | 208.49 | 0.915 | 119.92 | 30.91 | 203.3 | 0.956 |
| **HRM (2)** | | | | | | | | | | | | |
| Equal tpp | 200.70 | 31.85 | 206.7 | 0.991 | 201.55 | 18.79 | 226.0 | 0.993 | 134.93 | 23.88 | 272.8 | 0.806 |
| Equal dp | 195.13 | 25.11 | 204.1 | 0.961 | 201.98 | 18.59 | 225.4 | 0.975 | 126.27 | 13.95 | 266.3 | 0.078 |
| mbl | 209.92 | 33.69 | 206.1 | 0.971 | 228.83 | 19.80 | 222.3 | 0.868 | 187.92 | 18.36 | 267.0 | 0.964 |
| **HRM (3)** | | | | | | | | | | | | |
| Equal tpp | 204.85 | 31.77 | 214.4 | 0.997 | 193.65 | 19.71 | 261.2 | 0.994 | 145.45 | 21.76 | 335.0 | 0.982 |
| Equal dp | 200.48 | 24.98 | 212.3 | 0.978 | 213.46 | 19.36 | 255.8 | 0.694 | 108.11 | 14.99 | 324.7 | 0.998 |
| mbl | 205.96 | 33.88 | 214.8 | 0.985 | 227.41 | 28.34 | 252.2 | 0.748 | 141.30 | 27.53 | 328.8 | 0.559 |

TABLE S4. Statistics of the 63 model/tree combinations tested. Abbreviations: Info. = Information; Uncert.=Uncertainty; Avem.= estimated likelihood of scales present at the avemetatarsalian node; tpp=timePaleoPhy; dp=DatePhylo.

| | Equal tpp | | Equal dp | | mbl | |
|---|---|---|---|---|---|---|
| | AIC | Mean error | AIC | Mean error | AIC | Mean error |
| ER UNORD | 224.4 | 0.399 | 240.3 | 0.419 | 227.9 | 0.391 |
| ER ORD | 213.7 | 0.433 | 245.3 | 0.443 | 205.1 | 0.414 |
| ER ED | 214.8 | 0.417 | 234.3 | 0.445 | 220.5 | 0.378 |
| SYM UNORD | 201.8 | 0.357 | 214.4 | 0.356 | 201.5 | 0.328 |
| SYM ORD | 191.0 | 0.368 | 206.2 | 0.380 | 190.3 | 0.345 |
| SYM ED | 211.4 | 0.397 | 228.4 | 0.417 | 220.7 | 0.370 |
| ARD UNORD | 219.6 | 0.357 | 226.1 | 0.330 | 218.8 | 0.341 |
| ARD ORD | 184.3 | 0.361 | 188.5 | 0.347 | 181.6 | 0.343 |
| ARD ED | 196.2 | 0.382 | 211.2 | 0.394 | 200.1 | 0.387 |
| ER SMM-ind | 208.8 | 0.422 | 225.2 | 0.449 | 212.3 | 0.390 |
| ER SMM-sw | 211.0 | 0.425 | 227.6 | 0.459 | 214.5 | 0.402 |
| SYM SMM-ind | 201.8 | 0.410 | 216.0 | 0.431 | 208.49 | 0.415 |
| SYM SMM-sw | 206.8 | 0.423 | 217.7 | 0.456 | 212.5 | 0.463 |
| ARD SMM-ind | 206.1 | 0.433 | 205.3 | 0.455 | 203.3 | 0.417 |
| ARD SMM-sw | 206.4 | 0.437 | 205.3 | 0.456 | 203.5 | 0.420 |
| ER HRM (2) | 206.7 | 0.390 | 204.1 | 0.388 | 206.1 | 0.369 |
| ER HRM (3) | 214.4 | 0.379 | 212.3 | 0.388 | 214.8 | 0.367 |
| SYM HRM (2) | 226.0 | 0.370 | 225.4 | 0.377 | 222.3 | 0.348 |
| SYM HRM (3) | 261.2 | 0.377 | 255.8 | 0.361 | 252.2 | 0.346 |
| ARD HRM (2) | 272.8 | 0.371 | 266.3 | 0.328 | 267 | 0.343 |
| ARD HRM (3) | 335.0 | 0.366 | 324.7 | 0.318 | 328.8 | 0.344 |

TABLE S5. Mean error calculated for each of the tree-model combinations using a leave-one-out cross-validation (LOOCV) approach.
